# The CUP-SHAPED COTYLEDON 2 - MIR164A regulatory feedback loop directs leaf morphogenesis

**DOI:** 10.64898/2026.10.07.757447

**Authors:** Sophie Bigot, Aude Maugarny, Nia Verdon, Aurélie Vialette, Magali Goussot, François Roudier, Jasmine Burguet, Patrick Laufs, Nicolas Arnaud

**Author notes:** Correspondence (N. Arnaud).

## Abstract

Gene regulatory networks (GRNs) establish the spatio-temporal gene expression patterns necessary for reproducible development of organisms. MicroRNAs have emerged as important regulators of gene expression controlling either target gene levels, spatial patterns or timing. These molecular functions may vary depending on how they are integrated into GRNs. Here we used the Arabidopsis leaf as a model to study the genetic and molecular interactions between the CUP-SHAPED COTYLEDON 2 (CUC2) transcription factor and its negative regulator, miR164A. Quantitative morphometric analyses showed that miR164A acts from early stages during leaf development regulating *CUC2* expression dynamic rather than the spatial distribution of CUC2. At the molecular level, CUC2 directly activates *MIR164A* expression, forming a single negative feedback loop. Further, using a ratiometric approach, we identified a *cis*-regulatory element (CRE) sufficient to drive CUC2-dependent *MIR164A* expression at the leaf sinus. Ordinary differential equation (ODE) modelling suggests that CUC2-CRE may control the timely accumulation of CUC2. Accordingly, quantification of CUC2 protein accumulation dynamics using a spatio-temporal mean cartography strategy in CRISPR *cis*-regulatory variants at the *MIR164A* locus revealed that CUC2-dependent *MIRA164A* expression is required to maintain both the precise timing of *CUC2* expression and CUC2 levels. Altogether, our work provides experimental evidence for a specific role of the *CUC2-MIR164A* module in shaping leaf development, highlighting the contribution of network topology to GRN functions.

## Introduction

Multicellular organism development depends on the coordinated action of gene products within a dynamic and environment-dependent regulatory landscape. Gene expression programs are controlled by interconnected gene regulatory networks (GRNs) which implement non-linear regulatory logic through combinatorial control and regulatory loops (1–4). At the network level, transcriptional gene circuits are constituted of recurrent network motifs with specific functions *e.g.*, autoregulatory behaviors, feedback and feedforward loops, single-input modules (reviewed in (5)). GRNs can also involve regulators from distinct nature such as transcription factors (TFs), microRNAs or chromatin remodelers. For instance, motifs composed of TF-miRNA feedbacks are overrepresented in *Caenorhabditis elegans* and may confer robustness or oscillatory behaviors (6). Although the general function of network motifs has been extensively studied (5,7–10), their specific contribution to organismal development depends on how they are integrated into developmental frameworks.

Leaves initiate at the periphery of the shoot apical meristem as small cell clusters that progressively acquire their shape through cell proliferation, cell growth and differentiation. During these morphogenetic processes, the *Arabidopsis thaliana* leaf grows from a few micrometers to a few centimeters in length, progressively acquiring its shape and flat structure. The CUP-SHAPED COTYLEDON (CUC) transcription factors control serrations at the leaf margins and contribute to the leaf shaping. CUCs are members of the NO APICAL MERISTEM, ATAF2, CUP-SHAPED COTYLEDON (NAC) family conserved within the Green Lineage (11,12). There are three *CUC* genes in Arabidopsis (*CUC1-3*). In Arabidopsis, *CUC1* is not expressed in the developing leaf and loss of *CUC1* function does not affect leaf shape (13,14). CUC2 and CUC3 play partially redundant roles during serration development with CUC2 being required for teeth initiation and CUC3 for teeth growth maintenance (13). CUC2 acts as a trigger through molecular relays as a pulse of *CUC2* expression is sufficient to produce leaves with serrations in *cuc2* mutant background (15). However, CUC2 is not acting like a toggle switch, but more like a rheostat as plants with increasing CUC2 levels show deeper serrations at their leaf margins (15). Further, *CUC1* and *CUC2* are regulated by a microRNA, miR164, which is encoded by three different genes with distinct expression patterns and functions (*MIR164A, B* and *C*) (16,17). miR164 binding promotes *CUC2* mRNA cleavage (16) suggesting that it regulates *CUC* genes at the post-transcriptional level. Within this complex context, the leaf is a relatively simple structure, with *CUC2* and *MIR164A* genes acting as major contributors of leaf shape. Due to the complexity of interactions within GRNs, mathematical modeling approaches have been widely used to investigate how miRNAs shape the dynamics of gene regulatory networks (18). Indeed, features such as functional redundancy, modularity, and regulatory loops make genetic approaches challenging to design and interpret. Here we focus on the developing leaf of Arabidopsis as a tractable model system to study spatio-temporal gene expression dynamics at the organ level. Indeed, during leaf development, leaf shape acquisition requires only a subset of the *MIR164-CUC* components. Easy to access and image, the leaf is a good model to study genetic and molecular interactions between TFs and miRNA. Here, we first finely characterized the function of miR164 throughout leaf development. At the molecular level, we showed that CUC2 directly regulates *MIR164A* expression thus forming a single negative feedback loop. We identified a *cis*-regulatory element (CRE) with functional CUC2 binding sites which is sufficient to promote *MIR164A* expression at the leaf margin. Finally, using CRISPR-designed *cis*-regulatory *MIR164A* variants, we showed that this regulatory motif is pivotal for CUC2 function during leaf morphogenesis controlling the dynamics of CUC2 protein accumulation. Our work revealed that subtle changes in gene expression dynamics contribute to morphogenesis and showed how such TF-miRNA network motifs play instrumental roles in gene expression dynamics through their integration in developmental frameworks.

## Results and Discussion

### MIR164A controls leaf serration development from early stages

miR164 functions are partly dependent on its expression patterns (17). Among the three genes encoding miR164, *MIR164A* is the major contributor to leaf morphogenesis (14). For clarity, the miRNA produced from the *MIR164A* gene will hereafter be referred to as miR164A. The *mir164a-4* loss-of-function mutant shows increased serration levels, and it was suggested that miR164A might affect early stages of leaf development (14). However, this premise was not based on any precise morphological quantification. We conducted a thorough morphometric analysis of the *mir164a-4* leaves using the Morpholeaf software (19,20). First, rosettes and mean representations of mature leaf contours confirmed that the *mir164a-4* mutant displays more serrated leaves (Fig 1AB). This results partly from a small increase in serration number without changes in global leaf length (Fig 1C, Fig S1B). Second, *mir164a-4* teeth are sharper than the wild-type, mostly due to an increase in tooth height (Fig 1DE, Fig S1A, Fig S1C). This result applies to all tooth ranks along the leaf margin suggesting a general role for miR164A in controlling leaf serration outgrowth (Fig 1DE). To reveal the developmental origin of the phenotype observed in mature leaves, we analyze *mir164a-4* mutant leaves at early developmental stage, from initiation to 1-mm-long leaf primordia. Increased serration on the *mir164a-4* mutant can already be detected on a 250 μm long primordium (Fig 1F). Interestingly, teeth initiation rate was not different in *mir164a-4* compared to wild-type throughout young leaf development (Fig 1G), suggesting that the increase in teeth number observed on mature leaves may result from an extended period of tooth formation. At the tooth level, significant differences in tooth shape can be detected at early developmental stages (Fig1 IK), with *mir164a-4* teeth showing a strong increase in height for a small decrease in width compared to the wild-type (Fig 1HJ, Fig S1DE), consistent with observations on mature leaves. Overall, our morphometric phenotyping confirmed that miR164A negatively regulates serration outgrowth from early stages of leaf development.

**Figure 1:**
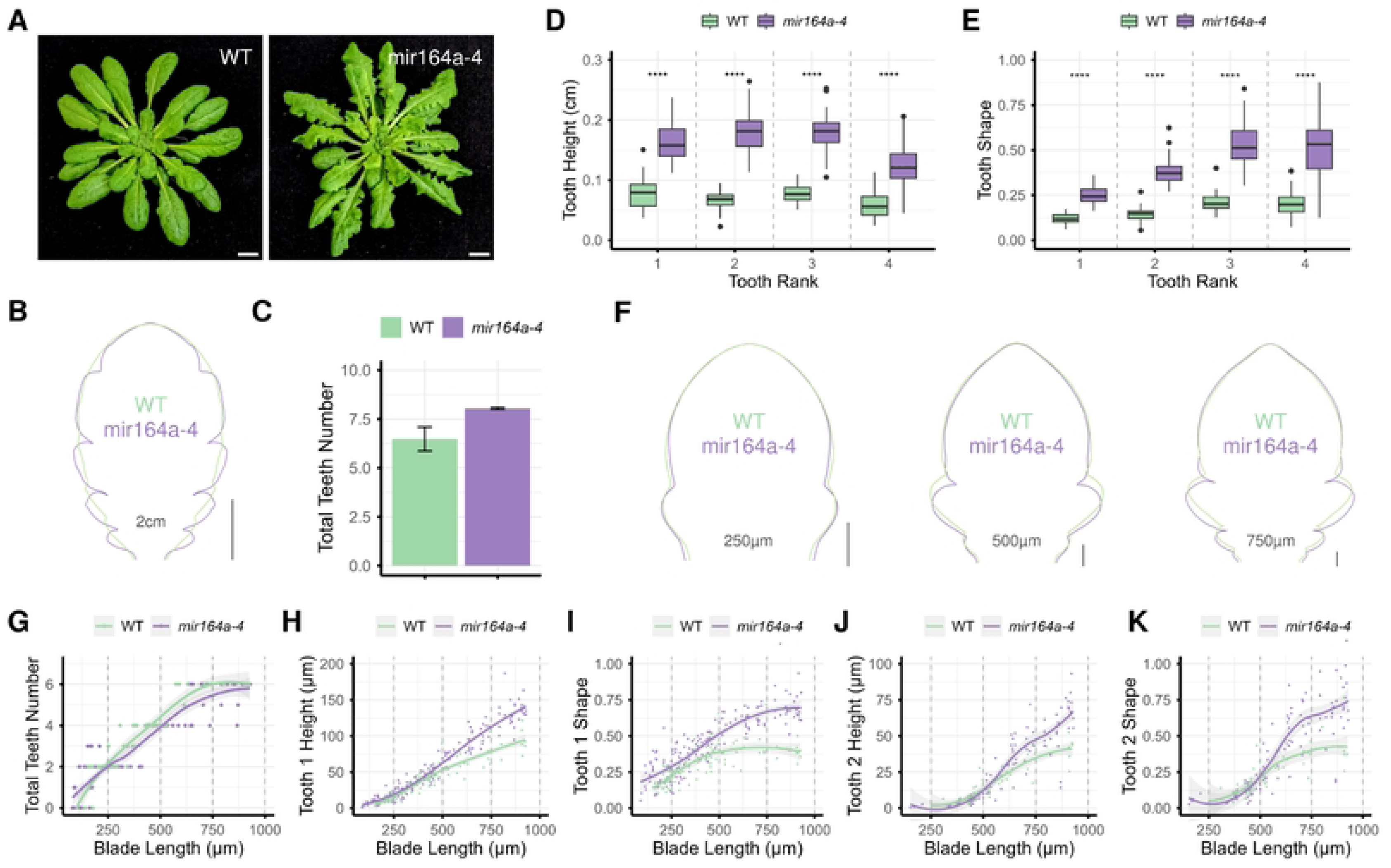
miR164A controls leaf shape from early developmental stages. (A) Rosettes of WT and *mir164a-4* mutant grown in short-day conditions for 6 weeks. Scale bar = 1cm. (B-E) Mean leaf silhouettes (B), total teeth number per leaf (C), tooth height (D) and tooth shape (E) of WT and *mir164a-4* mutant. Tooth shape is defined as the ratio between tooth height and tooth width as described in Fig S1A. Scale bar = 0.5 cm (B). (F) Mean silhouettes of WT and *mir164a-4* mutant developing leaves for three lengths (250µm, 500µm and 750µm). Scale bar = 50µm. (G) Teeth number along blade length in WT and *mir164a-4* mutant. (H-K) Tooth height (H and J) and tooth shape (I and K) evolution of WT and *mir164a-4* mutant along the leaf blade for respectively tooth 1 (H-I) and tooth 2 (J-K). (D-E) Statistical significance was assessed using a Wilcoxon test. Significance is indicated as follows: * p < 0.05, ** p < 0.01, *** p < 0.001, **** p < 0.0001. Leaves rank 11, 12 and 13 and tooth rank 1, 2, 3 and 4 from those leaves were used for leaf morphometrics.

### MIR164A controls CUC2 dynamics

miR164A negatively regulates *CUC2* expression during tooth development (14). In particular, we aimed to assess whether miR164 regulates CUC2 protein levels, timing and/or spatial distribution. First, we compared *CUC2* and *MIR164A* promoter activities at the leaf level using *pCUC2::erRFP* and *pMIR164A::erRFP* reporter lines: both reporters are expressed in sinuses during leaf development (Fig 2A). *MIR164A* promoter activity is also detected in the apical hydathode at the leaf tip (Fig 2A) as previously described in (14). To compare the localization of *CUC2* and *MIR164A* promoter activities at the cellular level, we imaged a double *pCUC2::erRFP* and *pMIR164A::GFP-N7* reporter line. A subset of sinus cells are marked with both erRFP and GFP-N7 signals suggesting that *CUC2* and *MIR164A* can be expressed simultaneously in sinus cells (Fig S2A). Other miRNA/TF regulatory modules playing important roles during leaf development, such as during leaf adaxial-abaxial polarity specification where HD-ZIPIII PHABULOSA (PHB) and PHAVOLUTA (PHV) transcription factors accumulate with a complementary pattern to that of miR165/miR166 (21). While this example illustrates how miRNAs establish distinct expression domains, *MIR164A* co-expression with its target in sinus cells suggests a different mode of action.

**Figure 2:**
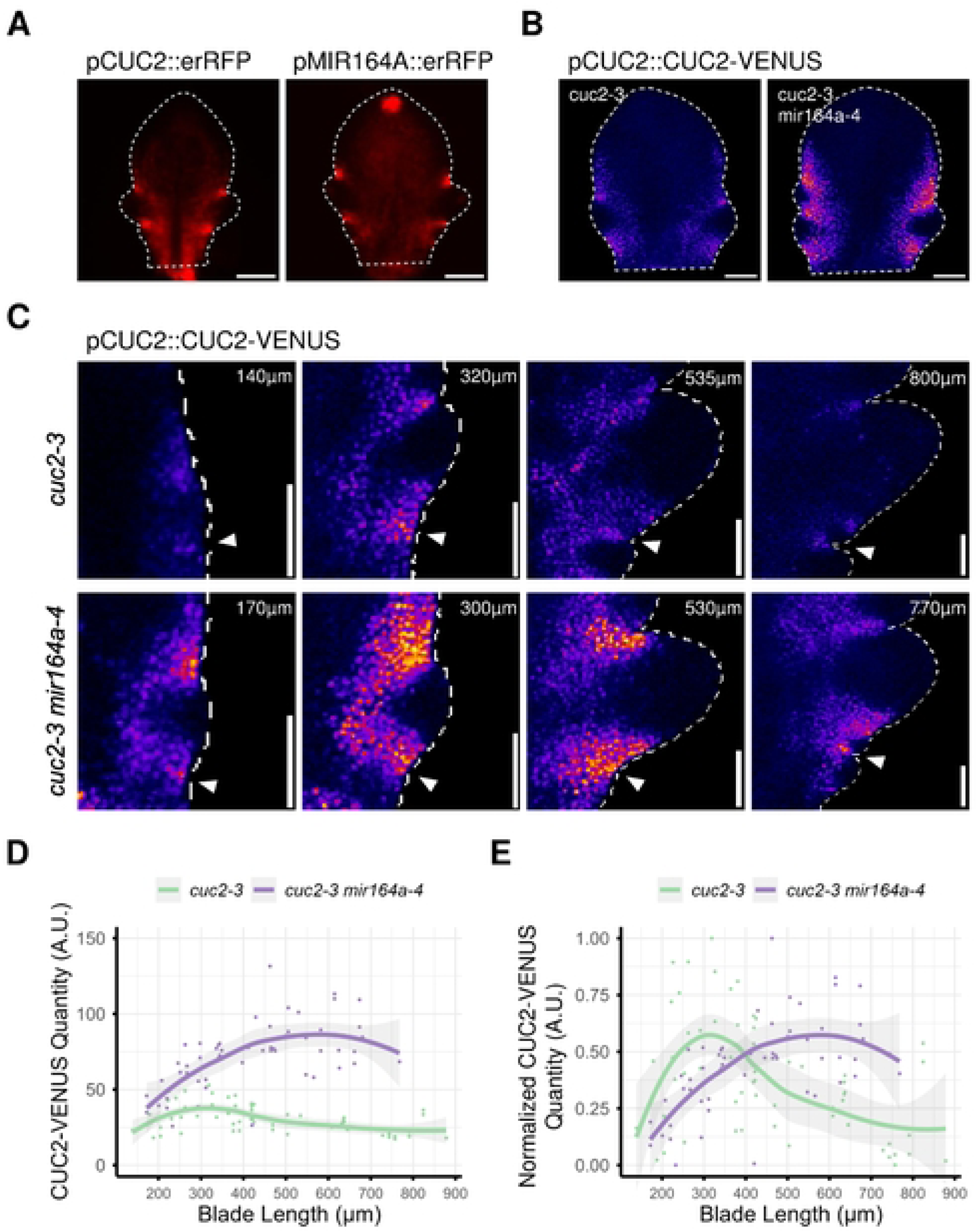
miR164A controls both levels and dynamics of CUC2 during early leaf development. (A) *pCUC2::erRFP* (left panel) and *pMIR164A::erRFP* (right panel) in developing leaves. Scale bar = 100µm. (B) *pCUC2::CUC2-VENUS* in *cuc2-3* (left panel) and in *cuc2-3 mir164a-4* (right panel) in developing leaves. Scale bar = 50µm. (C) *pCUC2::CUC2-VENUS* in *cuc2-3* and in *cuc2-3 mir164a-4* during early leaf development. Leaf length, used as a time proxy, is indicated for each image. These images are representative of the CUC2-VENUS dynamics measured throughout early leaf development in D and E. White arrowheads indicate the second sinus from where quantifications are realized in figures D and E. Scale bar = 50µm. (D-E) CUC2-VENUS signal intensity in sinus 2 for leaves of rank > 11 at early stages of leaf development in *cuc2-3* and in *cuc2-3 mir164a-4* (D), and normalized according to the minimum and maximum values for each line to compare CUC2-VENUS signal dynamics rather than levels (E).

To evaluate miR164A function, we examined *pCUC2::CUC2-VENUS* expression in either the *cuc2-3* or the *cuc2-3 mir164a-4* mutant backgrounds. Despite a clear increase in CUC2-VENUS level, whole-leaf observations do not reveal any major changes in CUC2-VENUS overall spatial distribution in the *cuc2-3 mir164a-4* background (Fig 2B). Indeed, CUC2-VENUS signal is still mainly found in the sinuses between emergent teeth, in a crescent shape around the outgrowth and excluded from the teeth. This pattern is reminiscent of the *pCUC2::erRFP* and pMIR164A::erRFP lines and we do not observe any discrepancies between *CUC2* transcriptional activity and CUC2 protein accumulation. This suggests that miR164A does not affect CUC2 spatial distribution. As the CUC2 spatial pattern is not changed in a *mir164a* mutant, the role of miR164A could be to dampen CUC2 levels and/or to modulate its accumulation timing.

To further examine this, we quantified CUC2-VENUS signal dynamics in the second sinus during early stages of leaf development using leaf length as a proxy of leaf age (Fig 2CDE). Our analysis revealed that, in *cuc2-3*, CUC2-VENUS is transiently expressed at early stages with a maximum accumulation in sinus 2 when the leaf is ∼300 µm long (Fig 2E). In the *cuc2-3 mir164a-4* mutant background, overall CUC2-VENUS levels increased ∼2.5 times (Fig 2D; Fig S2B), reaching a maximum when the leaf is ∼500 µm long with a broader dynamic (Fig 2E). Therefore, miR164A has a dual function: to dampen CUC2 levels and to fine-tune CUC2 protein accumulation dynamics at the leaf margin. This echoes a previous report showing that transient CUC2 activation with an inducible system in *cuc2* mutant is sufficient to trigger tooth initiation and that tooth development is subsequently maintained even in the absence of CUC2 (15). As CUC2 quantitatively regulates serration outgrowth, this miR164A-dependent regulation level may contribute to leaf morphogenesis.

### CUC2 directly activates MIR164A expression

Our quantitative data showed that *CUC2* and *MIR164A* are both expressed in leaf boundary domain (BD) cells. Moreover, previous data suggested that *MIR164A* expression may partly depend on CUC2 activity as *MIR164A* expression is altered specifically at the leaf margin in *cuc2-1* mutant background (13). It is then possible that *CUC2* and *MIR164A* are engaged in reciprocal interaction in part through the direct activation of *MIR164A* expression by CUC2. Such regulatory modules are indeed common within GRN and often contribute to peaks and/or oscillatory expression behaviors (6,7).

To test whether CUC2 can directly and positively control *MIR164A* expression, we first evaluated whether *MIR164A* expression depends on CUC2 levels. We analyzed the expression of the *pMIR164A::erRFP* reporter in young leaves from various genetic backgrounds with increasing levels of CUC2 *ie cuc2-3*, *WT*, *mir164a-4* and *CUC2g-m4*, a line expressing a miR164-resistant version of *CUC2* (Fig 3A). *MIR164A* promoter activity increased with increasing CUC2 levels, showing that CUC2 quantitatively controls *MIR164A* expression level. It is important to note that, despite the increase in signal intensity, the spatial expression pattern of *MIR164A* remains unchanged. *MIR164A* continues to be expressed in sinuses, where *CUC2* is also expressed (Fig 3A). *MIR164A* expression in the apical hydathode at the leaf tip does not vary with increasing CUC2 level suggesting that CUC2-dependent expression of *MIR164A* is leaf-margin specific (Fig 3A). Second, to test if CUC2 can activate *MIR164A* expression, we used a heterologous one-hybrid system coupled with flow cytometry assays, where *Physcomitrium patens* protoplasts were transfected with a *pMIR164A::GFP* construct in presence or absence of the CUC2 protein (22). Protoplasts transfected with *pMIR164A::GFP* alone showed a low level of background fluorescence (Fig 3B). When transfected together with a CUC2 expressing construct, a significant increase of the GFP signal intensity was detected (Fig 3B) suggesting that the CUC2 transcription factor is able to activate *MIR164A* promoter activity *in planta*. Third, in order to examine whether this CUC2-dependent *MIR164A* activation may also occur in Arabidopsis, we use a Dexamethasone (DEX)-inducible version of CUC2 (*cuc2-3 35S::CUC2-GR*) (23). Upon 6-hours DEX induction on plantlets, *MIR164A pre-miR* levels were significantly higher than in the mock-treated control showing that CUC2 induction is sufficient to promote *MIR164A* expression (Fig 3C). Fourth, we independently confirmed that CUC2 is able to activate the expression of *MIR164A* in F1 progeny between *35S::CUC2-GR* and two different *MIR164A* transcriptional reporters (*pMIR164A::erRFP* and *pMIR164A::GFP-N7*) upon DEX induction in root and leaf contexts, respectively (Fig S3A). Finally, to test whether this activation relies on direct interaction of CUC2 onto *MIR164A* promoter, we used Cycloheximide (CHX) as a translation inhibitor to reveal immediate targets for which translation is not required (24). DEX-induced CUC2-dependent accumulation of *pre-miR164A* transcript was resistant to CHX treatment (Fig 3D), suggesting that CUC2 directly regulates *MIR164A* expression.

**Figure 3:**
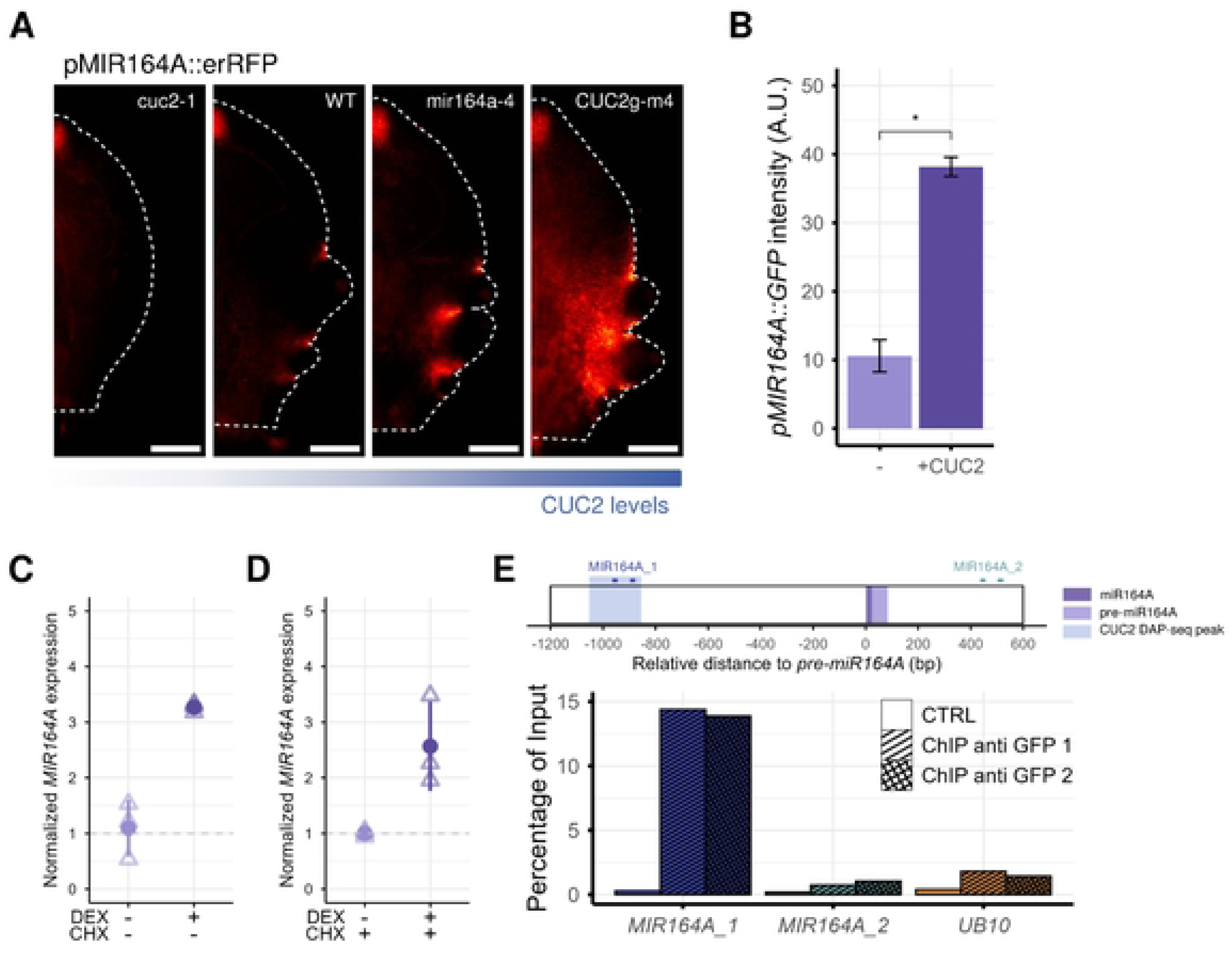
CUC2 activates directly *MIR164A* expression. (A) *pMIR164A::RFP* signal in genetic backgrounds with increasing CUC2 levels : *cuc2-1*, WT, *mir164a-4* and *CUC2g-m4*. Scale bar = 100µm. (B) *pMIR164A::GFP* signal in *Physcomitrium patens* protoplasts co-transfected or not with CUC2 CDS. (C-D) *MIR164A* expression level – measured as the *pre-miR* accumulation level – in a c*uc2-3 35S::CUC2-GR* line treated with either mock or dexamethasone (DEX) for 6h in the absence (C) or presence of cycloheximide (CHX) (D). (E) Chromatin immunoprecipitation (ChIP) experiment followed by qPCR on inflorescence apices from *pCUC2::CUC2-VENUS cuc2-3 mir164a-4* and *mir164a-4* used as a negative control sample, showing the direct interaction of CUC2-VENUS with the *MIR164A* locus. Relative enrichment is shown as percentage of input measured with primer pairs within *MIR164A* promoter region *(MIR164A_1)*, downstream of the *pre-miR164A* sequence (*MIR164A_2)* or for *UB10* as negative control gene (*UB10)*. Two biological replicates were performed for the test experiment (ChIP anti GFP 1 and 2, from *pCUC2::CUC2-VENUS cuc2-3 mir164a-4*) and one for the control condition (CTRL, from *mir164a-4* background). (C) Statistical significance was assessed using a Wilcoxon test. Significance is indicated as follows: * p < 0.05, ** p < 0.01, *** p < 0.001, **** p < 0.0001.

Accordingly, available DAP-seq data (25) predicted that CUC2 can bind to the *MIR164A* locus between -1053b to -854bp upstream from *pre-miR164A* (Fig S3B). Consistently, we identified four putative CUC2 binding sites (BS) within this DAP-seq peak based on the NAC TF core binding motif CGT[GA] defined in (26) (Fig S3B, Fig 3E and Fig 4A). Furthermore, phylogenetic shadowing revealed that, despite being intergenic, the region containing the putative CUC2 BS is highly conserved within the *Brassicaceae* family (Fig S3B). To unravel whether CUC2 can physically bind to *MIR164A* promoter region *in vivo*, we performed ChIP-qPCR assay in plants expressing CUC2-VENUS using a GFP antibody (Fig 3E). We performed the experiment on inflorescence apices of a line expressing *pCUC2::CUC2-VENUS* in a *cuc2-3 mir164a-4* mutant background in which older buds and mature flowers were removed. Indeed, *CUC2* and *MIR164A* are also expressed in flower BD and using floral buds of the *mir164a-4* mutant unable us to enriched ChIP tissue samples in BD cells with increased CUC2 level. For enrichment analysis, we targeted the upstream region defined by DAP-seq data, putative CUC2 BS and conservation assays (*MIR164A_1* primer pair) as well as a region located downstream *pre-miR164A,* as a negative control (*MIR164A_2* primer pair). In two biological replicates, a clear enrichment was observed in plant expressing CUC2-VENUS using *MIR164A_1* primers, but not with *MIR164A_2* primers or control primers (*UB10*), indicating that CUC2-VENUS binds to *MIR164A_1* promoter region *in planta*. Together, our data indicated that CUC2 directly activates the expression of *MIR164A* thus forming a single negative feedback loop that may contribute to CUC2 function during leaf development.

**Figure 4:**
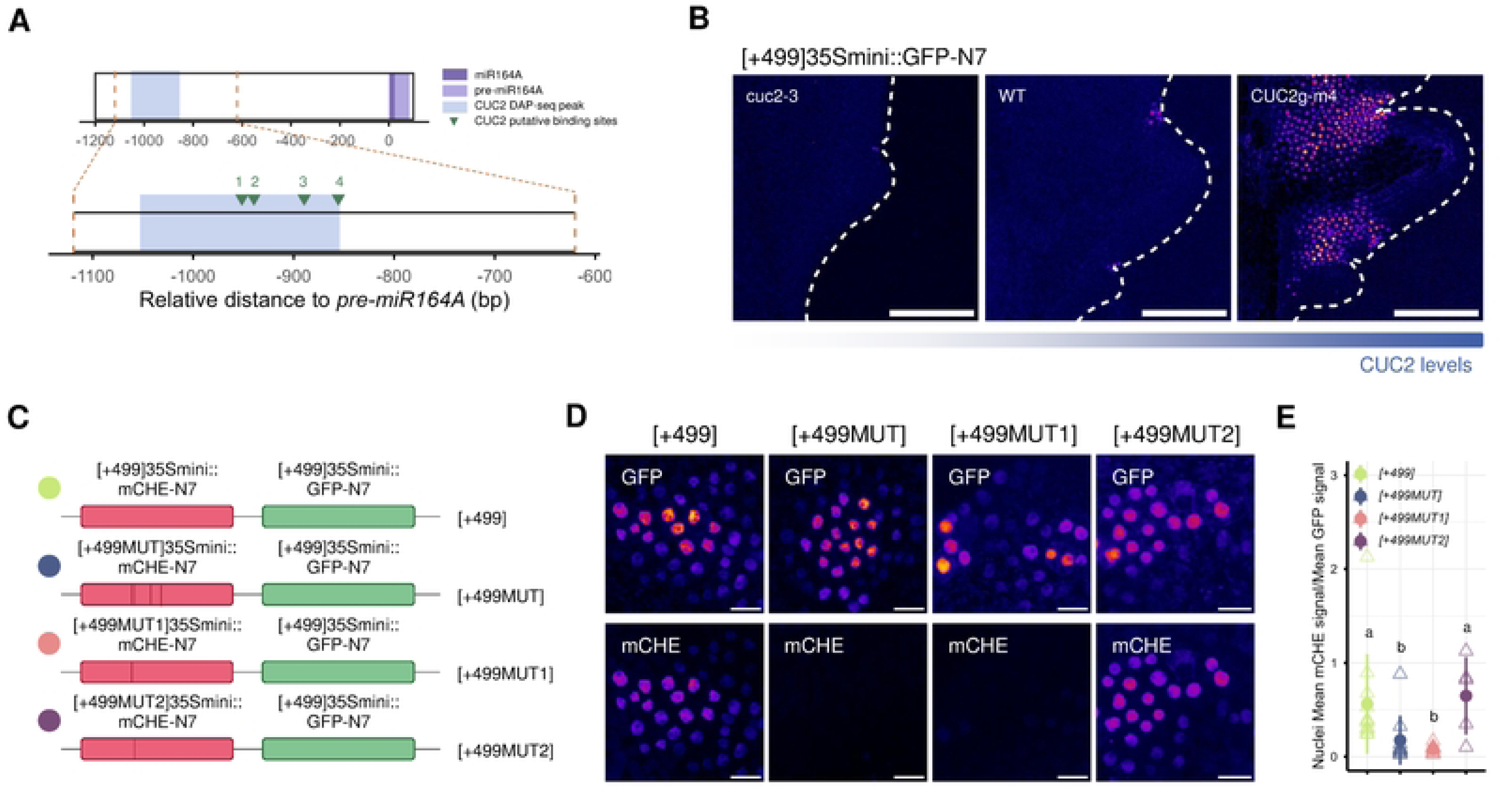
A CUC2-depend CRE is sufficient to promote *MIR164A* expression. (A) Scheme representing *MIR164A* locus with the position of putative CUC2 binding site (BS) indicated as green arrows 1-4. (B) *[+499]35Smini::GFP-N7* expression in genetic backgrounds with increasing CUC2 levels: *cuc2-1*, WT and *CUC2g-m4*. Scale bar = 100µm. (C) Scheme with the different ratiometric constructs used to analyze CUC2-dependent CRE. (D) Representative images from ratiometric transgenic lines for which GFP and mCHERRY signals, accounting for WT or mutated versions of CUC2-CRE respectively, were acquired simultaneously. Scale bar = 10µm. (E) Quantification of the mCHERRY over GFP fluorescence intensity ratio in independent T1 transgenic lines (n=12 for *[+499]*, n=10 for *[+499MUT]*, n=10 for *[+499MUT1]* and n=5 for *[+499MUT2]*). Values represent the mean ratio per line derived from single-nucleus quantifications. (E) Statistical significance was assessed using a Kruskal-Wallis test followed by a Dunn’s test. p-values were adjusted with the Bonferroni method. Statistical differences between groups are indicated by different letters.

### Identification of CUC2-dependent activating CRE within MIR164A promoter

Based on our finding that CUC2 activates the expression of *MIR164A* by directly binding to its promoter region, we further characterized the *cis*-regulatory element (CRE) involved in the regulatory interaction (Fig 4A). First, to test whether the region bound by CUC2 within the *MIR164A* promoter is sufficient to promote sinus-located expression *in vivo*, we designed and produced a construct with a 499bp fragment of the *MIR164A* promoter encompassing CUC2-dependent CRE (CUC2-CRE) fused to a minimal *35S* promoter driving the expression of a nuclear GFP namely *[+499]35Smini::GFP-N7*. The minimal *35S* promoter has been described as having little to no transcriptional activity on its own, whereas the insertion of enhancers element upstream results in transcriptional activity, allowing to test enhancer functions (27). Indeed, no transcriptional activity was detected at the leaf margin for control lines expressing *35Smini::GFP-N7* (Fig S4A). Independent transgenic lines in WT background carrying *[+499]35Smini::GFP-N7* construct showed a highly localized expression in sinus epidermal cells (Fig 4B and Fig S4B), suggesting that the *[+499]* fragment is sufficient to drive expression at the leaf margin in developing tooth sinuses. This expression depended on CUC2 levels as the GFP signal intensity in the sinus was low in plants with reduced CUC2 levels (*cuc2-3),* whereas in plants with elevated CUC2 levels (*CUC2g-m4*), the GFP signal was higher and more extended than in the WT while remaining restricted to the sinuses (Fig 4B) as observed for *pMIR164A::erRFP* line (Fig 3A). Accordingly, using the DEX-inducible *CUC2* overexpressing line *35S::CUC2-GR*, *[+499]35Smini::GFP-N7* expression was activated in the root upon DEX addition showing that CUC2 is sufficient to promote synthetic reporter expression independently of its developmental context (Fig S4C). Hence, we observed a CUC2-dependent expression of *[+499]35Smini::GFP-N7,* suggesting that CUC2 acts through its interaction with the CUC2-CRE to regulate *MIR164A* expression.

We further tested the roles of the individual CUC2 BS identified within CUC2-CRE using a ratiometric system derived from *[+499]35Smini::GFP-N7*. Briefly, we designed several modular constructs harboring two transcriptional units: *[+499]35Smini::GFP-N7* as an internal control, and *[+499]35Smini::mCHERRY-N7* or *[+499(mut)]35Smini::mCHERRY-N7* variants with BS substitutions (CGT[GA] → TTT[GA]) within CUC2-CRE (Fig 4C). For *[+499(mut)]*, we generated three different transcriptional units: two carrying substitutions in the distal CUC2 BS (BS1 and BS2) at positions -953 bp (*[+499MUT1]*) and -941 bp (*[+499MUT2]*) relative to *pre-miR164A*; and one (*[+499MUT]*) in which all four identified CUC2 binding sites were mutated, including BS1 and BS2. Because both GFP and mCHERRY reporters are cloned in a single vector, the nuclear mCHERRY/GFP ratio in individual transgenic plants enabled quantitative assessment of transcriptional activity changes induced by BS motif substitutions. Using this method, directly in T1 progeny, we observed a strong reduction in mCHERRY signal relative to GFP in the *[+499MUT]* lines compared to *[+499]* in the sinus (Fig 4D). Quantification revealed a significant decrease in the nuclear mCHERRY/GFP mean signal ratio (Fig 4E). Similar results were observed in *[+499MUT1]* lines, whereas no difference was detected in *[+499MUT2]* lines compare to *[+499]* (Fig 4DE). This suggests that *[+499MUT]* and *[+499MUT1]* drive gene expression less efficiently than *[+499]* and *[+499MUT2]*. Therefore, BS1 is the main contributor to the *MIR164A* expression activity promoted by CUC2 binding *in vivo* in this gain-of-function context. Our ratiometric approach enabled us to identify a functional CUC2 BS that accounts for the majority of CUC2-CRE expression capacity.

### MIR164A cis-regulatory variants disrupt CUC2 dynamics

Based on the identification and our characterization of CUC2-CRE within the *MIR164A* promoter, we investigated the effect of *cis*-regulatory mutations *in planta* at the *MIR164A locus*. To do so, we employed a CRISPR/Cas9 strategy to induce deletions within the region previously identified as bound by CUC2. We designed several RNA guides targeting this region and obtained two *cis*-regulatory mutants: *mir164a-Δ116* and *mir164a-Δ483* (Fig 5A). The *mir164a-Δ116* allele corresponded to a 116 bp deletion affecting the first three identified CUC2 BS, BS1 to BS3. The *mir164a-Δ483* allele contained a 483bp deletion together with several substitutions encompassing the entire CUC2-CRE, thereby affecting all four predicted CUC2 BS. Based on gain- of-function analysis of CUC2-CRE (Fig4 CDE), CUC2 binding in *MIR164A* promoter is likely affected in these two *cis*-regulatory mutants.

**Figure 5.**
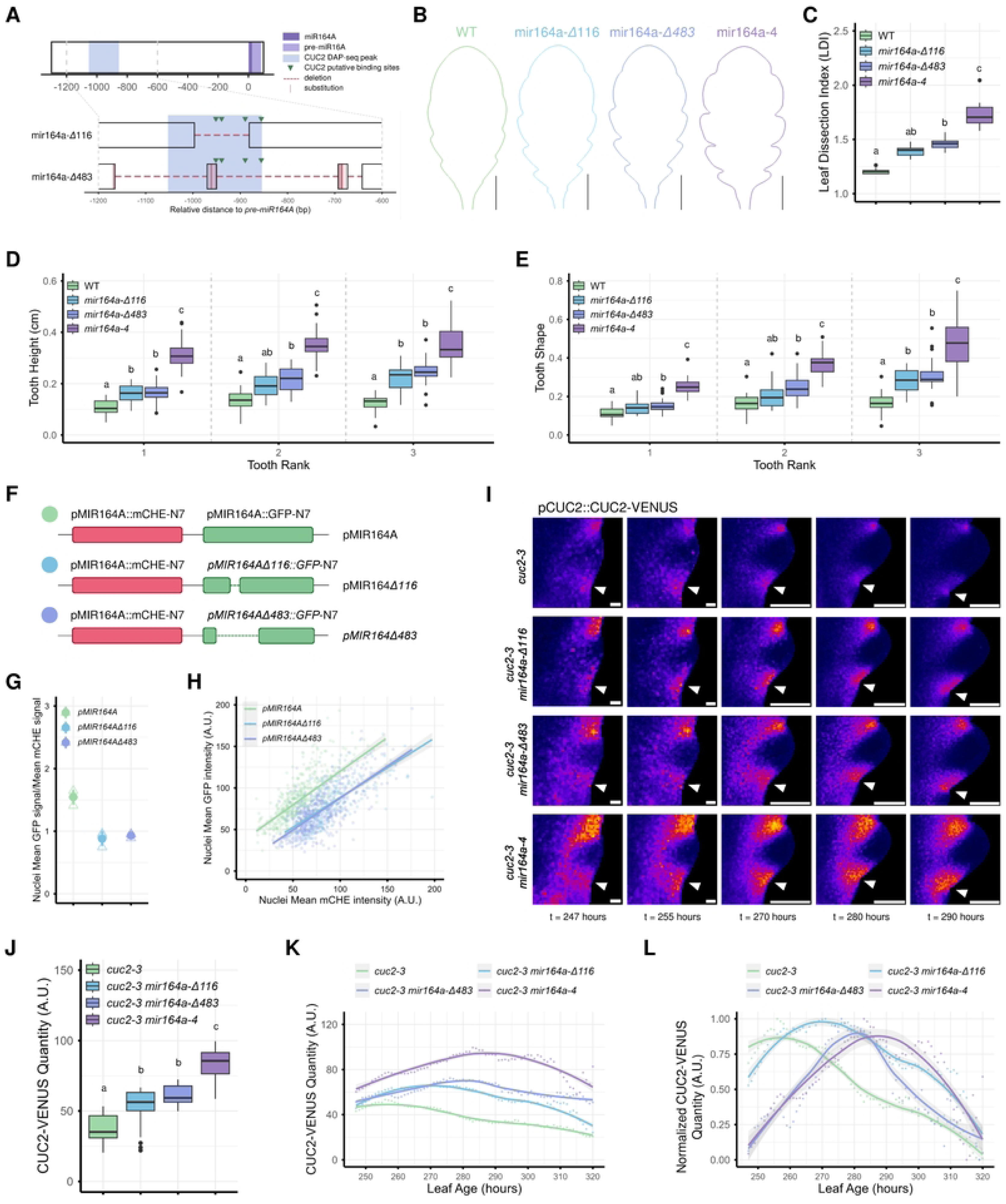
*Cis*-regulatory variants of *MIR164A* disrupt CUC2 dynamics. (A) Scheme representing *MIR164A* locus in the two independent *cis*-regulatory mutants, *mir164-Δ116* and *mir164-Δ483*. The positions of putative CUC2 binding site are indicated by green arrowheads. (B) Mean silhouettes of mature leaf shapes of WT, *mir164-Δ116*, *mir164-Δ483* and *mir164a-4* mutants. Scale bar = 1cm; n=13 for WT, n=10 for *mir164-Δ116,* n=13 for *mir164-Δ*483 and n=14 for *mir164a-4*. (C) Leaf Dissection Index (LDI) from leaves 11, 12 and 13 of WT, *mir164-Δ116*, *mir164-Δ483* and *mir164a-4.* (D-E) Tooth height (D) and tooth shape (E) of WT, *mir164-Δ116*, *mir164-Δ483* and *mir164a-4* from leaves rank 11, 12 and 13 and tooth rank 1, 2, 3. (F) Scheme with the different ratiometric constructs used to analyze *MIR164A* promoter activity of WT and *cis*-regulatory variants. (G) Quantification of the GFP over mCHE fluorescence intensity ratio in independent and monolocus T2 transgenic lines. Values represent the mean ratio per line (n=3) derived from single-nucleus quantifications). (H) Mean GFP intensity plotted against the mean mCHERRY intensity in three independent transgenic lines per genotype. Linear regressions were added for each genotype. (I) Representative images from the spatio-temporal mean cartography analysis showing CUC2-VENUS signal at different developmental time in lines expressing *pCUC2::CUC2-VENUS* in c*uc2-3, cuc2-3 mir164-Δ116, cuc2-3 mir164-Δ483* and *cuc2-3 mir164a-4*. Every single image is calculated from several individual images as a mean expression map for a given time point (see Material and Methods). Scale bars = 50µm. White arrows show the second sinus from where quantifications are realized in figures J-K-L. (J) CUC2-VENUS levels in developing leaves (leaf rank > 11), without size distinction, in c*uc2-3, cuc2-3 mir164-Δ116, cuc2-3 mir164-Δ483* and *cuc2-3 mir164a-4*. (K-L) Mean CUC2-VENUS signal intensity in sinus 2 for leaves of rank > 11 throughout leaf development in c*uc2-3, cuc2-3 mir164-Δ116, cuc2-3 mir164-Δ483* and *cuc2-3 mir164a-4* (K), and normalized for each line to compare CUC2-VENUS signal dynamics rather than levels (L). (C-D-E-J) Statistical significance was assessed using a Kruskal-Wallis test followed by a Dunn’s test. p-values were adjusted with the Bonferroni method. Statistical differences between groups are indicated by different letters.

We first analyzed the mature leaf shape phenotype of these mutants in comparison with WT and *mir164a-4* (Fig S5A). Mean representations of mature revealed increased margin dissection in the *cis*-regulatory mutants compared to WT, but less pronounced than in *mir164a-4* (Fig 5B), as confirmed by the Leaf Dissection Index (LDI) measurements (Fig 5C; (28)). This phenotype was associated with an increased serration number similar to *mir164a-4*, while leaf blade length remained unchanged (Fig S5BC). Quantitative analyses also revealed that the *cis*-regulatory mutants display sharper teeth (Fig 5E), with an intermediate phenotype between WT and *mir164a-4*. This was primarily driven by an increase in tooth height, with minimal contribution from changes in tooth width (Fig 5D, Fig S5D), similarly to what we previously observed in the *mir164a-4* mutant.

To assess the extent to which these *cis*-regulatory mutations affect *MIR164A* transcriptional activity *in vivo*, we used the same ratiometric approach as described in Fig 4CDE. We generated constructs harboring two transcriptional units: *pMIR164A::mCHE-N7* as an internal control, and either *pMIR164A::GFP-N7*, *pMIR164AΔ116::GFP-N7* or *pMIR164AΔ483::GFP-N7* (Fig 5F). The *pMIR164AΔ116* and *pMIR164AΔ483* correspond to the mutated promoter of *mir164a-Δ116* and *mir164a-Δ483 cis*-regulatory mutants respectively. As for the GFP signal, we observed that both mutated promoters remain active at the leaf margin, with an expression pattern still restricted to the sinus region, similar to the mCHERRY pattern (Fig S5E). This suggested that *cis*-regulatory mutations in *MIR164A* promoter do not affect the spatial expression pattern of *MIR164A*.

However, quantification revealed a significant decrease in the nuclear GFP/mCHERRY mean signal ratio in both mutated lines compared to the wild-type construct (Fig 5G). Similarly, an alternative representation of GFP signal intensity as a function of mCHERRY one showed that, for a given mCHERRY intensity, GFP signal was consistently lower in both mutated constructs (Fig 5H). Together, these results suggested that *MIR164A* promoter is overall less active in the *cis*-regulatory mutants *mir164a-Δ116* and *mir164a-*Δ483 than in WT. Interestingly, these *cis*-regulatory mutations did not lead to a complete loss of *MIR164A* expression, highlighting the involvement of additional *cis*-regulatory elements and their associated transcription factors in the regulation of *MIR164A* expression in the sinus.

To assess the functional relevance of CUC2 activation on *MIR164A* expression, we investigated the consequences of the *cis*-regulatory mutations on CUC2 accumulation. First, we first generated an ordinary differential equation (ODE) model consisting of 3 nodes: *CUC2* mRNA (*CUC2m*), CUC2 TF (CUC2p) and miR164A. Three coupled linear equations describe a simple negative feedback regulation mechanism of miR164A on the production of *CUC2* mRNA, an autoregulatory activation of CUC2 and an activation of miR164A expression dependent on CUC2 level (see schematic in Fig S6AB). The model is relative (non-dimensional) and the rate constants were selected such that the output matched the qualitative shape of the experimental curves for *pCUC2::CUC2-VENUS* dynamics in the *cuc2-3* background (Fig 2E). To test our ODE model, we simulate CUC2 behaviour in a *mir164a* mutant background. Suppression of the term corresponding to the miR164 regulation - without modifying other model parameters - lead to the loss of sharp regulatory dynamics of CUC2 (Fig 3C) as seen experimentally by monitoring CUC2-VENUS protein in a *cuc2-3 mir164a-4* mutant background (Fig 2EF). This shows that it is possible to reproduce CUC2-VENUS behavior with a simple three node model in at least two conditions including perturbations.

We then used our ODE model to simulate regulatory variations in *MIR164A* promoter by varying the parameter accounting for CUC2-dependent miR164A activation. In our model, decreasing the CUC2p activation of mir164A decreased the sharpness of the temporal regulation with a maximum of CUC2 protein accumulation delayed (Fig S5E) predicting that the direct activation of miR164A transcription by CUC2 within this FBL could promote the sharp and timely CUC2 accumulation. To confirm this prediction, we used the *pCUC2::CUC2-VENUS* line in the *cuc2-3 mir164a-Δ116* and *cuc2-3 mir164a-Δ483* backgrounds, in addition to the *cuc2-3* complemented background and the *cuc2-3 mir164a-4* one, similarly to what was described in Fig 2. In order to quantify CUC2-VENUS signal, we developed a quantitative approach allowing reconstruction of the signal mapped onto reference leaf shapes over a continuous ∼70-hour time course of leaf development. This method allows the quantification of fluorescent signal through time in a growing reference buffering variations due to measurement bias as we expected subtle changes in *cis*-regulatory mutants compare to the controls. Briefly, a large number of half-leaves from reporter lines were dated based on the blade length (between ∼150µm and ∼800µm, Fig S7A). The mean signal was computed in the reference morphospace for each time point (Fig S7BC). The contribution of each half-leaf in the average representation depended on its proximity to the target time point on the time axis. By repeating this process across multiple time points, we obtained a continuous overview of CUC2-VENUS dynamics. Similarly, to what was described for the *cuc2-3 mir164a-4* mutant, we did not observe any changes in the spatial distribution of CUC2-VENUS at the leaf margin throughout early leaf development in both *cis*-regulatory mutants, confirming that the CUC2-dependent expression of miR164A has no role in regulating CUC2 spatialization (Fig 5I). Quantification of CUC2-VENUS signal in sinus 2 of leaves between 247 and 320 hours revealed higher CUC2 levels in the *cuc2-3 mir164a-4* compared to *cuc2-3*, and intermediate levels with the *cis*-regulatory mutants (Fig 5J). Consistent with the results shown in Figure 2, analysis of CUC2 dynamics over time again revealed a transient accumulation of CUC2 in the complemented background, with a maximum around 255 hours, whereas this peak was delayed in *cuc2-3 mir164a-4*, reaching a maximum around 285-290 hours. As predicted by our modelling approach (Fig S6E), in the *cis*-regulatory mutants, CUC2-VENUS dynamics also remained transient but were delayed relative to the complemented background, with maxima around 270 hours and 280 hours for *cuc2-3 mir164a-Δ116* and *cuc2-3 mir164a-Δ483* respectively (Fig 5KL). Similar results were obtained when CUC2-VENUS signal was quantified using the same methodology as in Fig 2 (Fig S5GH). Taken together our quantitative analysis showed that the role of the CUC2-dependent regulation of *MIR164A* is to fine-tune transient CUC2 protein accumulation during leaf shape development.

## Conclusion

Here we show that the function of miR164A in the leaf is to control both CUC2 levels and CUC2 timing of accumulation. Furthermore, CUC2 directly activates *MIR164A* expression. In this case, coupling miRNA transcription to the activity of its target through a feedback loop (FBL) mechanism fine-tunes the target expression dynamics required for reproducible leaf development. This function is consistent with the finding that miR164A controls leaf shape from early stages of leaf development. To date, our understanding of miR164 function has been mostly obtained by the characterization of *mir164* mutants and miR164-resistant versions of its targets (14,16). Here we show that miR164A function is more complex than simply dampening CUC2 levels, highlighting the fact that molecular functions also need to be addressed at the network level.

Beyond providing molecular insights into the role of network topology during leaf development, our work also has evolutionary implications, highlighting for instance that subtle regulatory changes may impact developmental processes in different developmental contexts. For instance, during flower development, *MIR164C* expression is associated with boundary domain but not *MIR164B* (17), raising the possibility that CUC2 also regulates *MIR164C* expression. This could reflect evolutionary origins within the *MIR164* family where B-Clade *MIR164* genes (including *AthMIR164B*) and other *MIR164* genes (such as *AthMIR164A* and *AthMIR164C*) have diverged prior to the radiation of the Angiosperms (29). Further experiments will be required to investigate whether the *CUC2-MIR164A* module represents a general mechanism for modulating boundary domains activity and differential growth.

## Material and Methods

### Plant material and generation of transgenic plants

All genotypes are in the Columbia-0 (Col-0) background except the *cuc2-1* mutant originally isolated in Landsberg (Ler-0) back-crossed five times to Col-0 (13). The *CUC2g-m4* line and the *mir164a-4* mutant were previously described (14), as well as *pMIR164A::erRFP and pCUC2::erRFP* (15,30). *cuc2-3 pCUC2::CUC2-VENUS* and *mir164a-4 cuc2-3 pCUC2::CUC2-VENUS* lines were previously described in (15) and *cuc2-3 35S::CUC2-GR* in (23).

All cloning for the generation of transgenic plants was realized using GoldenBraid 2.0 technology (31). All vectors used are listed in Table S1. For the generation of *pMIR164A::GFP-N7,* the 2.1-kb promoter of *MIR164A* (14) was amplified on Col-0 genomic DNA using primers listed in Table S2 and inserted into a pUPD2 vector. *pMIR164A, GFP, N7* and *terminator 35S* parts were inserted into a pDGB3 α2 then combined with pDGB3 α1R binary vector containing *Tnos::NPTII::pNOS* into a pDGB3 Ω2.

For the generation *[+499]35Smini::GFP-N7*, a fragment of *MIR164A* promoter was amplified using primers listed in Table S2 and inserted in a pUPD2 vector. All parts were inserted into a pDGB3 α2 then combined with pDGB3 α1R containing *Tnos::NPTII::pNOS* into a pDGB3 Ω2. For the generation of *35Smini::GFP-N7*, pDGB3 Ω2 containing *Tnos::NPTII::pNOS/[+499]35Smini::GFP-N7* was digested with SpeI and XbaI to excise *[+499]* fragment prior self-ligation.

For the generation of *[+499]35Smini::mCherry-N7/[+499]35Smini::GFP-N7,* first, *[+499]35Smini::mCherry-N7* was assembled into a pDGB3 α2 and then into a pDGB3 Ω1 using dummy primers annealed before the reaction. This vector was combined with a pDGB3 Ω2 containing *Tnos::NPTII::pNOS*/*[+499]35Smini::GFP-N7* to generate the final construct. *[+499MUT], [+499MUT1]* and *[+499MUT2]* parts were first synthesized by Twist Bioscience (https://www.twistbioscience.com) and then assembled as above.

*mir164-Δ116* and *mir164-Δ483* mutants were generated for this study using CRISPR/CAS9 editing. Guide sequences designed using CRISPOR (https://crispor.gi.ucsc.edu/) are listed in Table S4. Primers for guide sequences were first annealed before being assembled with *pU6* and *sgRNA* into a pDGB3 α1 or α2. After successive combinations, the pDGB3 α1 containing all 4 transcriptional units was combined with a α2 containing *pCMV::DSRed:tnos/pRPS5a::hcas9:TRbcSE9* (kind gift from Lionel Gissot). Transgene was segregated in the T2 generation. Homozygous plants were genotyped using primers in Table S2.

For the generation of *pMIR164A::mCHE-N7/pMIR164A::GFP-N7,* first, *pMIR164A, mCHE, N7* and *terminator 35S* parts were inserted into a pDGB3 α2 and then in a pDGB3 Ω1 using dummy primers annealed before the reaction. This binary vector was then combined with previously described pDGB3 Ω2 *Tnos::NPTII::pNOS/pMIR164A::GFP-N7.* For the generation of *pMIR164A::mCHE-N7/pMIR164AΔ116::GFP-N7*, *pMIR164AΔ116* part was amplified on *mir164-Δ116* genomic DNA using primers listed in Table S2 and cloned in pUPD2. pDGB3 Ω2 *Tnos::NPTII::pNOS /pMIR164A::GFP-N7* and *pUPD2 pMIR164AΔ116* were digested with SpeI and XbaI to convert *pMIR164A* into *pMIR164AΔ116*. This vector was then combined with pDGB3 Ω1 containing *pMIR164A::mCHE-N7*. For *pMIR164A::mCHE-N7/pMIR164AΔ483::GFP-N7, pMIR164AΔ483* was synthetized by Twist Bioscience (https://www.twistbioscience.com). Parts were first inserted into pDGB3 α2, then combined with pDGB3 α1R binary vector containing *Tnos::NPTII::pNOS*. This vector was finally combined with pDGB3 Ω1 containing *pMIR164A::mCHE-N7*.

For the generation of all lines, wild-type plants were transformed through *Agrobacterium tumefaciens* floral dip method. For reporter lines generation, several lines with monolocus insertion with similar expression patterns were selected, one was chosen for imaging and crossing. Ratiometric analyses with gain of function constructs were directly conducted on kanamycin resistant primary transformants. For ratiometric analyses of *pMIR164A* promoter versions, three independent lines with monolocus insertion were selected and imaged.

### Growth conditions

For this study, all genotypes, except the one below, were grown under short-day (SD) conditions. Seeds were soaked in water and kept for two days in the dark at 4°C before being sown under SD conditions detailed in Table S3.

Seeds from *cuc2-3 35S::CUC2-GR* were surface sterilized, sowed in liquid Arabidopsis media (Duchefa S-Medium 11.8g/L, Ca(NO3)2 0.328g/L, MES 15mM, pH 5.8) and kept for two days in the dark at 4°C. Plants were grown under stirring at 120rpm and under long-day (LD) conditions [16 hours days (23,3°C, hygrometry 50%, 56μmol m-2 s-1 light), 8 hours night (23,3°C, hygrometry 50%)].

F1 seeds from the cross between *pMIR164A::GFP-N7* and *35S::CUC2-GR*, *pMIR164A::erRFP* and *35S::CUC2-GR, [+499]35Smini::GFP-N7* and *35S::CUC2-GR* were surface sterilized, sowed on 1% agar Arabidopsis media supplemented with MOCK (EtOH) or 10µM dexamethasone (DEX) (Duchefa S-Medium 11.8g/L, Ca(NO3)2 0.328g/L, MES 15mM, pH 5.8) and kept for two days in the dark at 4°C. Plants were grown vertically (for root imaging) or horizontally (for leaf imaging) in culture tower under long-day (LD) conditions [16 hours days (18°C, 50μE m-2 s-1, hygrometry not controlled), 8 hours night (18°C, hygrometry not controlled)].

For ratiometric approaches, seeds were surface sterilized, sowed on 1% agar Arabidopsis media supplemented with 50mg/L kanamycin (Duchefa S-Medium 11.8g/L, Ca(NO3)2 0.328g/L, MES 15mM, pH 5.8) and kept for two days in the dark at 4°C. Plants were grown horizontally in culture tower under long-day (LD) conditions [16 hours days (18°C, 50μE m-2 s-1, hygrometry not controlled), 8 hours night (18°C, hygrometry not controlled)]. 1-2-week-old plants resistant to kanamycin were then transferred in the soil under SD conditions (see Table S3).

For ChIP-qPCR experiment, *cuc2-3 mir164a pCUC2::CUC2-VENUS* and *mir164a-4* were grown in a greenhouse.

### Morphometric analyses

For morphometric analysis of mature leaves, leaves from ranks 11, 12 and 13 from 6 or 7-week-old plants were harvested and glued on a paper sheet prior to numerization. For morphometric analysis of developing leaves, young leaf primordia (ranks 11, 12 and 13) were dissected using a stereomicroscope throughout development starting 3-4 weeks after sowing. Leaves were mounted between a slide and a coverslip in a buffer containing 10 mM Tris HCl, pH 8.5, and 0.01% (v/v) Triton X-100 and imaged using an Axio Zoom.V16 microscope (Carl Zeiss Microscopy). Depending on the developmental stage imaged, either the chlorophyll fluorescent signal or the brightfield signal were collected.

Leaf silhouettes and quantifications were obtained using the MorphoLeaf software (19,20).

### Imaging

For *pCUC2::erRFP, pMIR164A::erRFP*, *cuc2-1 pMIR164A::erRFP*, *mir164a-4 pMIR164A::erRFP*, *CUC2g-m4 pMIR164A::erRFP*, leaves of rank 11, 12 and 13 from 4-week-old plants were dissected using a stereomicroscope. For the F1 *pMIR164A::erRFP x 35S::CUC2-GR*, leaves of rank 6 from 2-week-old plants were dissected. Leaves were mounted in a mounting medium (Tris-HCl 10mM, pH 8.5, Triton 0.01%) between a slide and a coverslip and imaged with an AxioZoom.V16 macroscope with an 80x zoom. A custom-made filter block (excitation band pass filter 560/25; beam splitter 585, emission band pass filter 615/24) was used to image RFP.

For leaf reporters analysis, leaves from rank 11 of 3-4 week old plants were dissected using a stereomicroscope. Leaves were mounted in a mounting medium (Tris-HCl 10mM, pH 8.5, Triton 0.01%) between a slide and a coverslip. For root reporters analysis, roots of 5-day-old plants were mounted in a mounting medium (Tris-HCl 10mM, pH 8.5, Triton 0.01%) between a slide and a coverslip. For *[+499]35Smini::GFP-N7*, 3-4 week-old-plants were fixed in 4% paraformaldehyde under a vacuum for 1h. Samples were then cleared in Clearsee (10% xylitol, 25% urea, and 15% deoxycholate), supplemented with SR2200 dye (32) for a week. Leaves from rank 11 were dissected using a stereomicroscope and mounted in a mounting medium (Tris-HCl 10mM, pH 8.5, Triton 0.01%) between a slide and a coverslip. All these samples were imaged on a Leica SP5 inverted microscope. The lenses were Leica 20x, 40x, 63x HCX PL APO CS using oil as an immersion medium. All acquisitions parameters are detailed in Table S5. Acquisition parameters were kept constant for experiments in which signal intensity was compared in different genetic backgrounds or when DEX was used.

All confocal microscopy images shown are maximum intensity projections generated in Fiji from acquired z-stacks, except for Figure S4B.

### Signal quantification

*pCUC2::CUC2-VENUS* signal was quantified in sinus 2 using Qpixies macro for ImageJ described in (30).

The quantification of ratios for gain-of-function experiments was performed as follows. In ImageJ, a circular region of interest (ROI) corresponding to each visible nucleus was manually traced based on the GFP channel. The “Measure” command available in ImageJ was then used to retrieve the mean intensity value for both GFP and mCherry channels. The same methodology was used for *pMIR164A::mCherry-N7/pMIR164A::GFP-N7, pMIR164A::mCherry-N7/pMIR164AΔ116::GFP-N7* and *pMIR164A::mCherry-N7/pMIR164AΔ483::GFP-N7*, this time using the mCherry channel to identify the nuclei. The ratio mCherry/GFP or GFP/mCherry ratio was then calculated for each nucleus using the mean intensities.

### *Physcomitrium patens* one-hybrid like system

All expression constructs were generated using the Gateway cloning technology (Invitrogen). *CUC2* CDS was previously cloned into the pDONR201 donor vector. The *CUC2* CDS fragment was cloned into the *pACTIN R1R2* destination vector *via* an LR Clonase recombination reaction to obtain *pACTIN::CUC2*. A fragment of the *MIR164A* promoter was amplified by PCR from Col-0 genomic DNA using pMIR164A_Forward and pMIR164A_Reverse primers (Suppl. Table 2), cloned into pDONR207 then subsequently transfered into the pBS R1R2 GFP destination vector to obtain p*BS pMIR164A::GFP.* Both vectors were used to transfect *Physcomitrium patens* protoplasts and the one-hybrid like plant system analysis was performed as described in (22).

### VISTA alignments

For *MIR164A* promoter alignments, the 1200 bp genomic sequence upstream of *pre-miR164A* of A. thaliana was aligned to *Arabidopsis lyrata*, *Arabidopsis halleri*, *Capsella rubella*, *Boechera stricta*, *Brassica rapa* subsp. *Pekinensis*, *Brassica rapa* subsp. *Rapa* and *Glycine max* genomes using VISTA-Point (VISTA Gateway (lbl.gov)). The conservation score was calculated on a 100-bp window with a minimum 90% conservation identity.

### Chromatin immunoprecipitation and qPCR

Chromatin immunoprecipitation was performed as described in Morao et al., 2016 with some modifications. Briefly, inflorescence apices from *pCUC2::CUC2-VENUS cuc2-3 mir164a-4* and control *mir164a-4* plants were hand dissected, snap-frozen in liquid nitrogen and ground. Double cross-linking was performed by treating grounded tissues with 2 mM di(N-succinimidyl) glutarate for 45 minutes at room temperature followed by 1% formaldehyde for 7 minutes. Cross-linked samples were then homogenized using a douncer. Chromatin was extracted from a nuclei-enriched pellet and extracted chromatin was sonicated using a Covaris S220 ultra-sonicator (12 min with 5% duty cycle, 105W peak power, and 200 cycles per burst) to generate DNA fragments with an average length of 250 bp. A small fraction of sonicated chromatin of each replicate was kept as input, while the rest was immunoprecipitated overnight with anti-GFP (Abcam, ab290) antibodies coupled to magnetic beads (Dynabeads, Invitrogen). DNA was recovered from both immunoprecipitated fraction and input by proteinase K treatment, reverse crosslinking at 65°C overnight and phenol/chloroform extraction followed by Qiagen Minelute column purification and concentration. 1 ng of immunoprecipitated DNA was used for library preparation using MicroPlex library preparation kit v2 (Diagenode). ChIP experiments were performed in at least two biological replicates. The optimal number of cycles needed to generate each library was estimated by real-time PCR.

Relative enrichment levels were determined by ChIP-qPCR in an optical 384-well plate in the QuantStudio 6 Flex Real-Time PCR System (ThermoFisher Scientific), using FastStart Universal SYBR Green Master (Rox) (Roche), according to the manufacturer’s instructions. The data were analyzed using the QuantStudio Real-Time PCR Software v1.3 (Applied Biosystems). Cp values were converted using the 2-ΔCT method and relative enrichment was expressed as percentage of input. qPCRs were performed in 2 technical replicates for each sample and at least two biological replicates were analyzed independently. Primers used for this assay are given in Table S2.

### ODE modeling

The model consists of 3 coupled linear ordinary differential equations, representing the dynamics of 3 variables, *CUC2m*, CUC2p protein and mirna164a over time (dt) (Fig S6A). These are represented by m, c and m164, respectively in the equations below:

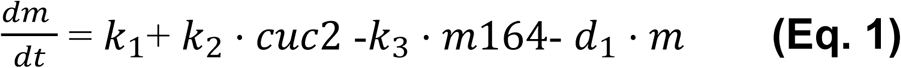

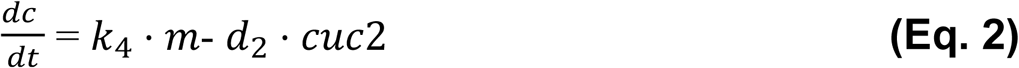

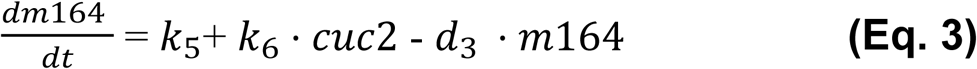

**Eq. 1** represents the CUC2m level, including both the auto-activation of CUC2 which positively influences the production of mRNA, and the repression of this process by miR164A.

**Eq. 2** represents the production of CUC2 protein.

**Eq. 3** represents the production of miR164A, including the activation of miR164A by CUC2p.

Both *CUC2m* and miR164A are produced at independent rates (k_1_ and k_5_, respectively) when there is no interaction with the other variables. Whereas CUC2p is only produced when there is *CUC2m*. All three equations have a degradation term, reducing the amount of the variable over time.

The model parameters were selected qualitatively, relative to the CUC2p production constant (k_4_), as it is not possible to extract individual network parameters from experimental expression intensities (which represent the output of the model).

Strong regulatory dynamics are only observed when the suppression constant (k_3_) is relatively high, which is expected as a microRNA can rapidly decrease mRNA levels. We found that to achieve tight regulatory dynamics, it was also required to set the rate constant for m164 smaller than the CUC2 activation factor (k_6_ > k_5_). This aligns with the findings that CUC2 is the main activator of miR164A (14).

A range of values was found which satisfied these requirements whilst also remaining non-negative at all time points. A set of default values for the parameters used for the simulations shown in this paper are given in Fig S6B.

All model results presented in the paper were calculated with the initial conditions at 0 for each variable. However, the regulation dynamics are robust against perturbations of the initial conditions (Fig S6D). For both the mutants studied, only the relevant parameter was altered relative to the WT. In the *mir164a* mutant, k_3_ = 0 (representing no suppression by mir164a). In cis-reg mir164a1, k_6_ = 0, representing no activation of mir164a by CUC2. Varying the k_6_ parameter to simulate *cis*-regulatory variants show that decreasing the CUC2p activation of *MIR164A* decreases the precision of the temporal regulation with a maximum of CUC2 protein accumulation delayed (Fig S6E). The equations were solved using the default configuration of scipy.integrate.odeint in python 3.10.12 and plotted with seaborn.

### Spatio-temporal mean cartography method

The dynamics of the fluorescent gene reporters were computed using the method described by (33) (Fig S7). The average contours showing the evolution of the half-leaf shape during growth were pre-calculated using MorphoLeaf software (19,20). These contours serve as a developmental reference space in which the average cartography is computed. First, the contours of individual leaf images are delineated using MorphoLeaf. Then, each leaf is assigned an age according to its length measurement, and the leaves are sorted along the time axis. Next, the half-leaves are processed separately. At a given time point (*), the half-leaf images of leaves close to that time point are spatially normalized and projected onto the corresponding average leaf contour in the developmental reference space. The normalized half-leaf images are averaged, with each image being weighted according to its distance from the selected time point. This results in the mean signal in the average contour in the reference space. Repeating this procedure along the time axis enables the continuous reconstruction of the fluorescent signal dynamics.

### Expression analyses

10-day-old, liquid-grown *cuc2-3 35S::CUC2-GR* plants were treated with dexamethasone (DEX) (10 μM) and/or cycloheximide (CHX) (10μM) 10 minutes prior DEX treatment, or MOCK (EtOH). Whole seedlings were collected 6 hours after treatment and immediately frozen in liquid nitrogen. Total RNAs were isolated using RNAeasy Plant Mini Kit (Qiagen) following manufacturer instructions. Reverse transcription was performed on a 2μg of total RNA using Thermo Scientific RevertAid H Minus Reverse Transcriptase at 42°C for 1 hour, followed by RNAse H treatment at 37°C for 20 minutes. Real time PCR was performed using the SsoAdvanced Universal SYBR® Green Supermix following manufacturer instructions on a Bio-Rad CFX machine. Statistical analysis were using the ΔΔCt method. Primers used for real time PCR are available in Table S2.

### Statistical analysis

Data analysis, statistics and plots were performed using R software (http://www.R-project.org/) and the graphics package ggplot2.

## Acknowledgments

We thank Lionel Gissot for the kind gift of the pCMV::DSRed:tnos/pRPS5a::hcas9:TRbcSE9 construct. We also thank Jean-Christophe Palauqui, Véronique Pautot and Alice Vayssières for discussions and the critical reading of the manuscript.

## Author contributions

Conceptualization : SB, AM, PL, JB, NA; Formal analysis : SB, AM, NV, AV, MG, FR, PL, JB, NA; Investigation : SB, AM, NV, AV, MG, FR, PL, JB, NA; Writing - original draft preparation : SB, NA; Writing - Review & Editing : SB, AM, NV, AV, MG, FR, PL, JB, NA; Visualization : SB, AM, NV, NA; Supervision : PL, JB, NA; Funding acquisition : JB, NA.

